## Supplementary material for "A fast and robust Bayesian nonparametric method for prediction of complex traits using summary statistics": Suppplementary Note

### Table of Contents


| | |
|---|---|
| SDPR Model | 2 |
| Modified Likelihood Function | 3 |
| Dirichlet Process Prior | 5 |
| Connection of the modified likelihood function with LDSC and SumHer | 7 |
| Partition of the reference LD matrix | 8 |
| MCMC Algorithm | 10 |
| Selection of phenotypes in UK biobank | 14 |
| Supplementary Table 2-10 | 15 |
| Supplementary Figure 2 | 18 |
| Supplementary Figure 3 | 19 |

### SDPR Model

Suppose GWAS summary statistics are derived based on $N$ individuals and $p$ genetic markers, the phenotypes and genotypes can be related through a multivariate linear model,

$$y = X\beta + \epsilon \tag{1}$$

where $y$ is an $N \times 1$ vector of phenotypes, $X$ is an $N \times p$ matrix of genotypes, and $\beta$ is an $p \times 1$ vector of effect sizes. We further assume, without loss of generality, that both $y$ and columns of $X$ have been standardized. GWAS summary statistics usually contain the per SNP effect size $\hat{\beta}$ directly obtained or well approximated through the marginal regression $\hat{\beta} = \frac{X^T y}{N}$. From this approximation, one can derive the commonly used likelihood function,

$$\hat{\beta}|\beta \sim N(R\beta, \frac{R}{N}) \tag{2}$$

where $R = \frac{X^T X}{N}$ is the reference LD matrix. Importantly, one should note that this equation assumes that each SNP has approximately the same sample size, which may not be true in the real setting. Furthermore, sometimes the correlation of marginal effect sizes in the summary statistics may not agree with equation (2), as shown in the table below. Failure to account for such discrepancy may cause convergence issues for SDPR.

| SNP | A1 | A2 | beta | se | p | N | r2 |
|---|---|---|---|---|---|---|---|
| rs1206549 | A | G | 0.0093 | 0.0065 | 0.15 | 197888 | 0.99 |
| rs712951 | A | G | -0.0037 | 0.0041 | 0.41 | 252571 | |

Supplementary Table 1. Illustration of the model misspecification issue using two SNPs in height GWAS summary statistics [1]. In 1000G EUR samples, rs1206549 and rs712951 are in strong LD. However, their effect sizes in GWAS summary statistics were in the opposite direction, whereas the likelihood function $\hat{\beta}|\beta \sim N(R\beta, \frac{R}{N})$ would expect these two SNPs to have similar effect sizes. The discrepancy observed here may be explained by the fact that the imputed sample sizes of the two SNPs were different.

### Modified Likelihood Function

We next derive the likelihood function when SNPs are typed on different individuals, motivated by the observation in the supplementary table 1.

Claim: If each SNP j is genotyped on $N_j$ individuals, define the matrix $H$ whose elements are $H_{ii} = \frac{1}{N_i}$ and $H_{ij} = \frac{N_{s,ij}}{N_i N_j}$ $(i \neq j)$, where $N_{s,ij}$ is the number of shared individuals genotyped for SNPs i and j. Then the likelihood function can be evaluated as

$$\hat{\beta}|\beta \sim N(R\beta, R \circ H) \quad (3)$$

where ° is the Hadamard product.

Proof: Let $S_j$ be the set of individuals on which SNP j is genotyped ($N_j = |S_j|$), then

$$E\left[\widehat{\beta_j}\middle|\beta\right] = E\left[\frac{X_j^T y}{Nj}\middle|\beta\right]$$

$$= \sum_{i \in S_j} \frac{X_{ji}}{N_j}\left(\sum_k X_{ik}\beta_k\right)$$

$$= \sum_{k} \left( \sum_{i \in S_j} \frac{X_{ji} X_{ik}}{N_j} \right) \beta_k$$

$$= \sum_{k} R_{jk} \beta_k$$

For $i \neq j$, we have

$$cov\left[\hat{\beta}_i, \hat{\beta}_j \mid \beta\right] = E\left[\frac{X_i^T \epsilon_i X_j^T \epsilon_j}{N_i N_j} \middle| \beta\right]$$

$$= \frac{1}{N_i N_j} E\left[X_i^T \epsilon_i \epsilon_j^T X_j\right]$$

$$= \frac{1}{N_i N_j} X_i^T E\left[\epsilon_i \epsilon_j^T\right] X_j$$

$$= \frac{1}{N_i N_j} \sum_{k \in S_i \cap S_j} X_{ik} X_{jk}$$

$$= \frac{R_{ij} N_{s,ij}}{N_i N_j}$$

It is trivial to check that when the sample size of all SNPs is same, the derived likelihood function is the same as equation (2). Furthermore, the correlation of marginal effect sizes in the GWAS summary statistics will be less than the correlation in the reference panel, depending on how many individuals are overlapped ($N_{s,ij}$) for two SNPs ($cor = \frac{R_{ij} N_{s,ij}}{\sqrt{N_i N_j}} \leq \frac{R_{ij} \min(N_i, N_j)}{\sqrt{N_i N_j}} \leq R_{ij}$). As an extreme example, for two SNPs that are in perfect LD ($R_{ij} = 1$), if they are genotyped on completely nonoverlapped individuals ($N_s = 0$), the correlation of their effect sizes in the summary statistics would be zero.

Evaluation of equation (3) requires the knowledge about the sample size and inclusion of each study for each SNP. For example, SNPs of GWAS summary statistics of lipid traits were genotyped on two arrays in two separate cohorts (GWAS chip: $N_1 \approx 95{,}000$; Metabochip: $N_2 \approx 94{,}000$) [2]. Based on this information, $N_{s,ij}$ is set to 0 if SNPs i and j were genotyped on different arrays, $N_1$ if SNP i was genotyped on GWAS chip and SNP j was genotyped on both arrays, and $N_2$ if SNP i was genotyped on Metabochip and SNP j was genotyped on both arrays. When this information is not available, we consider evaluating the likelihood function from the following distribution

$$\frac{\widehat{\beta}}{c}|\beta \sim N\left(R\beta, \frac{R + NaI}{N}\right) \tag{4}$$

where $R$ is the reference LD matrix, $c$ is a constant to correct for deflation of summary statistics if double genomic control was applied, and $a$ is a constant to shrink the covariance between two SNPs. For simplicity of notification, we will denote deflation-corrected marginal effect sizes as $\hat{\beta}$, i.e. $\hat{\beta} \coloneqq \hat{\beta}/c$. The correlation of marginal effect sizes of two SNPs is $\frac{R_{ij}}{1+Na}$ in equation (4), similar to the shrunk correlation $\frac{R_{ij}N_{s,ij}}{\sqrt{N_i N_j}}$ in equation (3).

### Dirichlet Process Prior

Like many Bayesian methods, we assume that the effect size of i^th^ SNP $\beta_i$, follows a normal distribution with mean 0 and variance $\sigma_\beta^2$. In contrast to methods assuming one particular parametric distribution, we consider placing a Dirichlet process prior on $\sigma_\beta^2$, i.e.

$$\beta_i \sim N(0, \sigma_\beta^2), \sigma_\beta^2 \sim DP(H_\beta, \alpha) \tag{5}$$

where $H_\beta$ is the base distribution and $\alpha$ is the concentration parameter controlling the shrinkage of the distribution on $\sigma^2$ toward $H$. To improve the mixing of MCMC and avoid the informativeness issue of inverse gamma distribution, we expand the parameter $\beta_i = \eta\gamma_i$ and assign the following prior [3]:

$$\begin{aligned} \eta &\sim N(0, A), \\ \gamma_i &\sim N(0, \sigma^2), \\ \sigma^2 &\sim DP(H, \alpha) \\ H &= IG(a_{0k}, b_{0k}) \end{aligned} \tag{6}$$

We note that the product $\eta\gamma_i$ in (6) corresponds to $\beta_i$ in (5), and $|\eta|\sigma$ in (6) corresponds to $\sigma_\beta$ in (5). We set $A = 10^6, a_{0k} = 0.5, b_{0k} = 0.5$ so that marginally the base distribution $H_\beta$ in equation (5) is approximately the square of uniform distribution on $[0, +\infty)$. (If $\eta \sim N(0, A), \sigma^2 \sim IG(0.5, 0.5)$, then $\frac{\eta}{\sqrt{A}}\sigma \sim Cauchy(0,1)$ and $p(|\eta|\sigma) \propto 1$ as $A \to \infty$ ).

Under the truncated stick-breaking representation of Dirichlet process, the full model can be rewritten as:

$$\hat{\beta}|\gamma, \eta \sim N\left(R\eta\gamma, \frac{R + aNI}{N}\right)$$

$$\eta \sim N(0, 10^6)$$

$$\gamma_j|\sigma_k^2, p_k \sim \sum_{k=1}^{M} p_k N(0, \sigma_k^2), k = 1, \dots, M, j = 1, \dots, p$$

$$p_k = V_k \prod_{m=1}^{1-k} (1 - V_m), \;\; V_k|\alpha \sim Beta(1, \alpha),$$

$$\sigma_k^2 \sim IG(0,5, 0.5), \alpha \sim Gamma(0.1,0.1)$$

We set $M$ to 1000 as default for our methods. To assess whether our choice of $M$ was a good approximation to the infinite stick-breaking process model, we counted the number of variance components to which SNPs were assigned. It turned out that SNPs were assigned to only 600 to 800 components in simulations and real data applications. After all, if the number of variance components is infinite, then some variance components will have no assignments of SNPs. Therefore, we believe that our choice of $M$ was sufficient to approximate the Dirichlet process.

### Connection of the modified likelihood function with LDSC and SumHer

If we assume that $\beta_j \sim N(0, \frac{h^2}{M})$ and the likelihood function (4), after integrating out $\beta$ we get

$$\hat{\beta} \sim N(0, \frac{c^2 h^2}{M} R^T R + c^2 \frac{R}{N} + c^2 aI)$$

$$E\left[N\widehat{\beta_J}^2\right] = c^2[\frac{N}{M}(\sum_k r_{jk}^2) h^2 + 1 + aN]$$

If $c = 1$ (no deflation of marginal effect sizes), then this is equivalent to LDSC where $1 + aN$ corresponds to the intercept term in LDSC [4]. The correction for deflation is similar to SumHer [5]. In practice, we found that it is safe to set $aN$ to .1 for the purpose of estimating effect sizes. If there is significant amount of deflation (e.g. intercept of LDSC is significantly less than 1), then we recommend to run SumHer to obtain the deflation correction factor $C$ and set $c$ to the square root of $C$ [5] (only observed for BMI in real data applications).

### Partition of the reference LD matrix

Partitioning the reference LD matrix into independent blocks allows the use of efficient collapsed Gibbs sampler and parallel sampling of the assignment vector. At present, ldtect is widely used for performing the partition task [6]. Ldtect works by computing the antidiagonal sum of the covariance matrix and applying the signal filtering approach to find the local minima in order to set the breakpoint. Originally developed to facilitate the interpretation of GWAS association signals, ldetect is not optimized for providing a precise partition for computation of PRS.

We used simulation data (Scenario 4) to assess the accuracy of LD blocks provided by ldtect. We generated marginal effect sizes and plotted them against the theoretical ones assuming the likelihood function $\hat{\beta}|\beta \sim N\left(R\beta, \frac{R}{N}\right)$. Unexpectedly, we found that the marginal effect sizes of some SNPs did not agree with the likelihood function (Supplementary Figure 1). These SNPs were from a region (Chr10: 33 Mb) where ldtect cut the entire block into two independent ones (bottom left and upper right as separated by the cross). The cut was incorrect as SNPs in the second block had significant amount of correlation with SNPs in the first block. Therefore, some SNPs in the second blocks would have non-zero marginal effect sizes if they were in LD with the causal SNP in the first block, whereas theoretically they would have zero effect sizes assuming two blocks were independent.

To solve this issue, we designed a simple algorithm to ensure that each SNP in one LD block does not have nonignorable correlation ($r^2 > 0.1$) with SNPs in other blocks. Assuming

SNPs are sorted based on their physical locations on the chromosome, for each SNP we recorded the index of the rightmost SNP with the nonignorable correlation using a sliding window. We then computed the cumulative maximum of the index along the list. We set the breakpoint at the SNP whose cumulative maximum index equals its original index. When applied to the example mentioned above, our algorithm did not cut the block and the marginal effect sizes were consistent with the theoretical ones (Supplementary Figure 1). Compared with ldtect, overall our algorithm produced similar number of blocks. However, the number of the blocks containing more than 1000 SNPs was larger for our algorithm.

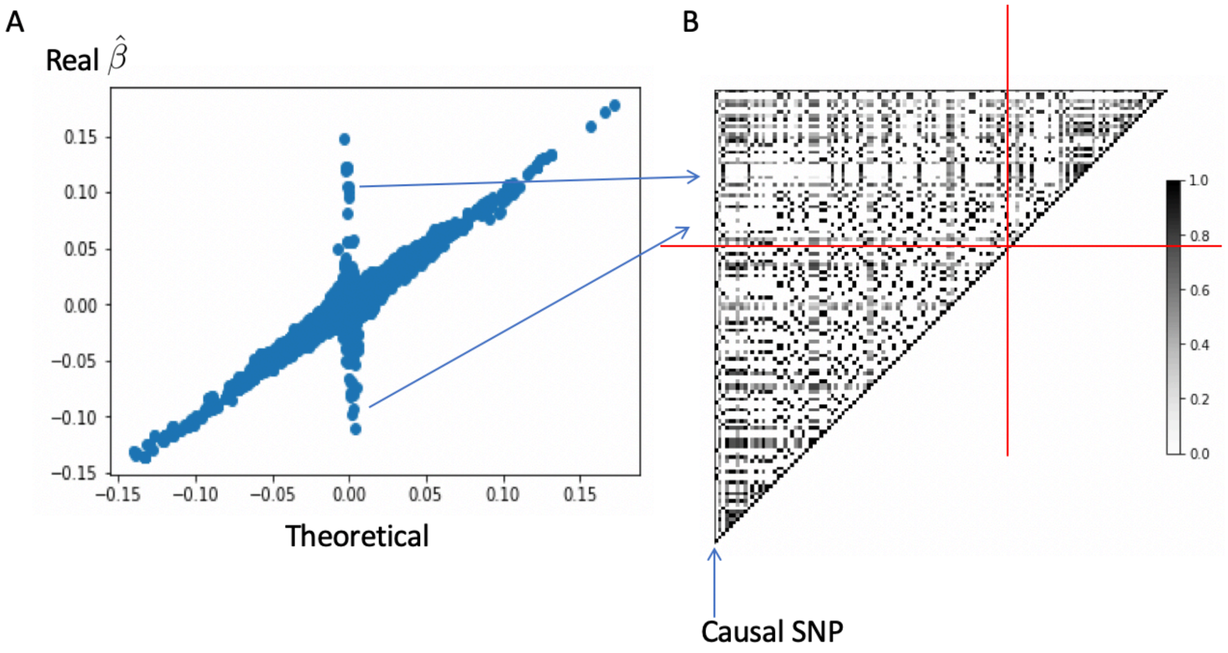

C

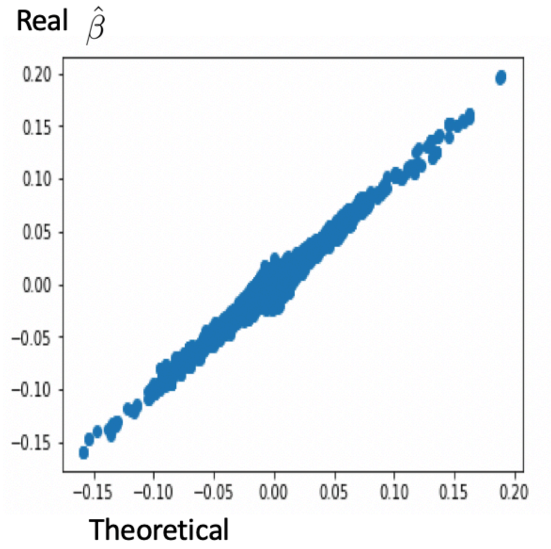


D

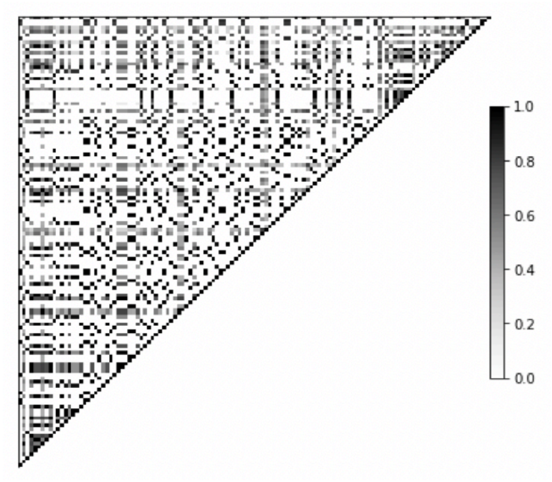


Supplementary Figure 1. (A) Comparison of theoretical (using the partition by ldtect) and marginal effect sizes in GWAS summary statistics. (B) Correlation matrix of SNPs in the Chr10:33 Mb region of UK Biobank genotype data. Ldtect divided these SNPs into two independent blocks as separated by the red cross. The upper-left dots indicated that SNPs in two blocks had nonignorable correlation and some SNPs in block 2 were in LD with the marked causal SNP. (C) Comparison of theoretical (using the partition by our method is correct) and marginal effect sizes in GWAS summary statistics. (D) Correlation matrix of SNPs in the Chr10:33 Mb region of UKB genotype data. Our method did not divide SNPs into two blocks.

### MCMC Algorithm

Here we describe our MCMC algorithm to obtain the posterior samples to estimate the effect sizes. We introduce a vector $z$ indicating the assignment of variance component for each SNP.

Compute $A, B$: $(R/N + aI)A = R$ or $(R \circ H)A = R$, $B = RA$. If $R \circ H$ is not positive definite, we add $-1.1 \times$ its minimum eigenvalue to the diagonal to make it positive definite.

Sampling $z_j$: For each LD block, we first integrate out the effect size $\gamma$ to derive the full conditional likelihood of $P\big(z_j = k \,\big|.\,\big)$:

$$P\big(z_j = k\big|.\big) \propto \int p\big(\hat{\beta}\big|\gamma,\eta\big)p\big(\gamma_j\big|z_j = k\big)\,d\gamma_j \;\times\; P\big(z_j = k\big)$$

$$\propto \int \exp\left\{-\frac{1}{2}\big(\hat{\beta} - \eta R\gamma\big)^T\left(\frac{R}{N} + aI\right)^{-1}\big(\hat{\beta} - \eta R\gamma\big)\right\}\frac{1}{\sigma_k}\exp\left\{-\frac{\gamma_j^2}{2\sigma_k^2}\right\}d\gamma_j \times p_k$$

$$\propto \int \exp\left\{-\frac{1}{2}\eta^2\gamma^T B\gamma + \eta\hat{\beta}^T A\gamma\right\}\frac{1}{\sigma_k}\exp\left\{-\frac{\gamma_j^2}{2\sigma_k^2}\right\}d\gamma_j \times p_k$$

$$\propto \int \exp\left\{-\frac{1}{2}\eta^2 B_{jj}\gamma_j^2 - \eta^2\sum_{i\neq j} B_{ij}\gamma_i\gamma_j + \eta\sum_i A_{ij}\hat{\beta}_i\gamma_j\right\}\frac{1}{\sigma_k}\exp\left\{-\frac{\gamma_j^2}{2\sigma_k^2}\right\}d\gamma_j \times p_k$$

$$\propto \int \exp\left\{-\frac{1}{2}\left(\eta^2 B_{jj} + \frac{1}{\sigma_k^2}\right)\gamma_j^2 + b_j\gamma_j\right\}d\gamma_j \;\times\frac{p_k}{\sigma_k}$$

$$\propto \frac{1}{\sqrt{\eta^2 B_{jj}\sigma_k^2 + 1}}\exp\left\{\frac{b_j^2}{2\big(\eta^2 B_{jj} + \sigma_k^{-2}\big)}\right\}p_k$$

where $b_j = \eta\sum_i A_{ij}\hat{\beta}_i - \eta^2\sum_{i\neq j} B_{ij}\gamma_i$. We set the first variance component to 0 in analogous to Bayesian variable selection, and we have $P\big(z_j = 1\big|.\big) \propto p_1$ as the integration equals 1 when $\gamma_j$ is degenerated at 0. We use log-exp-sum trick to avoid numerical overflow. Note that because SNPs in different LD blocks are approximately independent, we can sample their assignments in parallel.

Sampling $\beta$: We jointly sample the effect size of causal SNPs $\gamma_\theta$ in one independent LD block. The full conditional likelihood of $\gamma_\theta$ is

$$p(\gamma_\theta | z_j \neq 1, \ .) \propto \exp\left\{-\frac{1}{2}\eta^2\gamma^T B\gamma + \eta\hat{\beta}^T A\gamma\right\}\exp\left\{-\frac{1}{2}\gamma_\theta^T\Sigma_0^{-1}\gamma_\theta\right\}$$

$$\propto \exp\left\{-\frac{1}{2}\eta^2\gamma_\theta^T B_\theta\gamma_\theta + \eta\hat{\beta}^T A_\theta\gamma_\theta\right\}\exp\left\{-\frac{1}{2}\gamma_\theta^T\Sigma_0^{-1}\gamma_\theta\right\}$$

$$= MVN(\eta\Sigma A_\theta^T\hat{\beta}, \ \Sigma)$$

where $\Sigma = (\ \eta^2 B_\theta + \Sigma_0^{-1})^{-1}, \Sigma_0 = diag\left(\sigma_{z_1}^2, \dots, \sigma_{z_p}^2\right)$ for causal SNPs ($z_j \neq 1$). $A_\theta, B_\theta$ are the submatrices by selecting columns corresponding to SNPs with non-zero effect sizes from matrices $A, B$. For SNPs whose variance components are 0 ($z_j = 1$), we simply set the posterior effect sizes to 0 as $p(\gamma_j | z_j = 1, .) = 0$.

Sampling $\eta$: The full conditional likelihood is

$$p(\eta | .) \propto \exp\left\{-\frac{1}{2}\eta^2\Sigma\gamma_\theta^T B_\theta\gamma_\theta + \eta\Sigma\hat{\beta}^T A_\theta\gamma_\theta\right\}\exp\left\{-\frac{\eta^2}{2\times 10^{-6}}\right\}$$

$$= N\left(\frac{\Sigma\hat{\beta}^T A_\theta\gamma_\theta}{\sum\gamma_\theta^T B_\theta\gamma_\theta + 10^{-6}}, \ \frac{1}{\sum\gamma_\theta^T B_\theta\gamma_\theta + 10^{-6}}\right)$$

Sampling $\sigma_k^2$: The first variance component is always 0. The full conditional likelihood is

$$p(\sigma_k^2 | .) \propto \prod_{j:z_j=k}\frac{1}{\sigma_k}\exp\left\{-\frac{\gamma_j^2}{2\sigma_k^2}\right\}\sigma_k^{-2(a_{0k}-1)}\exp\left\{-\frac{b_{0k}}{\sigma_k^2}\right\}$$

$$= IG(\frac{M_k}{2} + .5, \frac{\sum_{j:z_j=k}\gamma_j^2}{2} + .5)$$

where $M_k = \sum_j I(z_j = k)$ and $I$ is the indicator function.

Sampling $V_k$: The full conditional likelihood is

$$p(V_k|.) \propto p_k^{M_k} \dots p_{M-1}^{M_{M-1}} p_M^{M_M} V_k^{1-1}(1-V_k)^{\alpha-1}$$

$$\propto V_k^{M_k}(1-V_k)^{M_{k+1}+\cdots+M_M+\alpha-1}$$

$$= Beta(1+M_k, \alpha+\sum_{l=k+1}^{M} M_l)$$

for k = 1, …, M – 1. $V_M$ equals 1 according to the definition of the truncated stick-breaking process.

Computing $p_k$: The prior probability can be computed as

$$p_1 = V_1$$

$$p_k = \prod_{l=1}^{k-1}(1-V_l)\, V_k \quad (k \geq 2)$$

Sampling $\alpha$: The full conditional probability is

$$p(\alpha|.) \propto \prod_{l=1}^{M-1} \alpha(1-V_l)^{\alpha-1}\alpha^{.1-1}\exp\{-.1\times\alpha\}$$

$$= Gamma(0.1+M-1, 0.1-\sum_{k=1}^{M-1}\log(1-V_k))$$

We record the effect size $\beta = \eta\gamma$ and heritability $h^2 = \beta^T R\beta$ for each iteration and compute the average of all posterior samples as the final estimator.

### Selection of phenotypes in UK biobank

We selected individuals with six quantitative traits based on relevant data fields (50 for height, 21001 for BMI, 30780 for LDL, 20760 for HDL, 20690 for total cholesterol, and 30870 for triglyceride). We used the first instance if multiple measurements were available. For six diseases, cases were selected based on ICD code in the EHR and self-reported questionnaire (data field 20002). For coronary artery disease, cases were selected based on ICD-9 codes of 410.X, 411.0, 412.X, or 429.79 or ICD-10 codes of I21.X, I22.X, I23.X, I25.2, or self-reported myocardial infarction [7]. For breast cancer, cases were selected among female participants based on ICD-9 codes 174 or 174.9, or ICD-10 codes C50.X, or self-report history of breast cancer. For inflammatory bowel disease, cases were selected based on ICD-10 codes of K50.X, or ICD-9 codes of 555.X, or self-reported history of Crohn's disease, ulcerative colitis, and inflammatory bowel disease. Participants with self-reported history of immunological/system disorders were excluded from controls. For type 2 diabetes, cases were selected based on ICD-10 codes of E11.X, or ICD-9 codes of K51.X, or self-reported history of type 2 diabetes. Participants with self-reported history of diabetes were excluded from controls. For schizophrenia, cases were selected based on ICD-10 codes of F20.X, or ICD-9 codes of 295.X, or self-reported history of schizophrenia. Participants with self-reported history of neurobiology/eye/psychiatry disorders were excluded from controls. For bipolar, cases were selected based on ICD-10 codes of F31.X, or ICD-9 codes of 296.X, or self-reported history of type I and type II bipolar disorder. Participants with self-reported history of neurobiology/eye/psychiatry disorders were excluded from controls.

### Supplementary Table 2-10

| Sample size | SDPR | PRS-CS | SBayesR | LDpred | P+T | LDpred2 | Lassosum | DBSLMM |
|---|---|---|---|---|---|---|---|---|
| 10K | 0.461 | 0.393 | 0.459 | 0.448 | 0.405 | 0.458 | 0.410 | 0.423 |
| 50K | 0.493 | 0.428 | 0.495 | 0.337 | 0.421 | 0.489 | 0.446 | 0.412 |
| 100K | 0.494 | 0.424 | 0.497 | 0.328 | 0.384 | 0.495 | 0.423 | 0.397 |

Supplementary Table 2. The median of square of Pearson correlation across 10 simulations for Scenario 1A.

| Sample size | SDPR | PRS-CS | SBayesR | LDpred | P+T | LDpred2 | Lassosum | DBSLMM |
|---|---|---|---|---|---|---|---|---|
| 10K | 0.200 | 0.137 | 0.209 | 0.209 | 0.168 | 0.217 | 0.186 | 0.168 |
| 50K | 0.426 | 0.354 | 0.424 | 0.332 | 0.358 | 0.426 | 0.381 | 0.364 |
| 100K | 0.462 | 0.405 | 0.459 | 0.289 | 0.378 | 0.463 | 0.425 | 0.386 |

Supplementary Table 3. The median of square of Pearson correlation across 10 simulations for Scenario 1B.

| Sample size | SDPR | PRS-CS | SBayesR | LDpred | P+T | LDpred2 | Lassosum | DBSLMM |
|---|---|---|---|---|---|---|---|---|
| 10K | 0.056 | 0.052 | 0.053 | 0.056 | 0.043 | 0.056 | 0.050 | 0.052 |
| 50K | 0.198 | 0.179 | 0.2 | 0.179 | 0.147 | 0.208 | 0.181 | 0.170 |
| 100K | 0.293 | 0.254 | 0.289 | 0.278 | 0.209 | 0.305 | 0.270 | 0.259 |

Supplementary Table 4. The median of square of Pearson correlation across 10 simulations for Scenario 1C.

| Sample size | SDPR | PRS-CS | SBayesR | LDpred | P+T | LDpred2 | Lassosum | DBSLMM |
|---|---|---|---|---|---|---|---|---|
| 10K | 0.385 | 0.335 | 0.386 | 0.375 | 0.345 | 0.383 | 0.348 | 0.355 |
| 50K | 0.449 | 0.386 | 0.447 | 0.343 | 0.376 | 0.449 | 0.396 | 0.382 |
| 100K | 0.462 | 0.389 | 0.461 | 0.311 | 0.363 | 0.463 | 0.399 | 0.375 |

Supplementary Table 5. The median of square of Pearson correlation across 10 simulations for Scenario 4.

| Sample size | SDPR | PRS-CS | SBayesR | LDpred | P+T | LDpred2 | Lassosum | DBSLMM |
|---|---|---|---|---|---|---|---|---|
| 10K | 0.050 | 0.046 | 0.048 | 0.054 | 0.042 | 0.054 | 0.050 | 0.053 |
| 50K | 0.157 | 0.146 | 0.146 | 0.157 | 0.133 | 0.159 | 0.151 | 0.150 |
| 100K | 0.215 | 0.204 | 0.197 | 0.210 | 0.177 | 0.216 | 0.207 | 0.205 |

Supplementary Table 6. The median of square of Pearson correlation across 10 simulations for Scenario 5.

| Traits | SDPR | PRS-CS | SBayesR | LDpred | P+T | LDpred2 | Lassosum | DBSLMM |
|---|---|---|---|---|---|---|---|---|
| Height | 0.271 | 0.271 | 0.226 | 0.23 | 0.210 | 0.267 | 0.269 | 0.261 |
| BMI | 0.093 | 0.089 | 0.082 | 0.088 | 0.076 | 0.093 | 0.092 | 0.082 |
| HDL | 0.116 | 0.114 | 0.033 | 0.096 | 0.079 | 0.114 | 0.106 | 0.104 |
| LDL | 0.147 | 0.141 | 0.026 | 0.133 | 0.105 | 0.143 | 0.142 | 0.120 |
| Total cholesterol | 0.146 | 0.144 | 0.032 | 0.139 | 0.111 | 0.145 | 0.141 | 0.129 |
| Triglycerides | 0.075 | 0.082 | 0.021 | 0.071 | 0.057 | 0.081 | 0.076 | 0.072 |

Supplementary Table 7. The mean of variance of phenotypes explained by PRS across 10 random splits for six quantitative traits.

| Traits | SDPR | PRS-CS | SBayesR | LDpred | P+T | LDpred2 | Lassosum | DBSLMM |
|---|---|---|---|---|---|---|---|---|
| CAD | 0.591 | 0.596 | 0.579 | 0.604 | 0.584 | 0.604 | 0.594 | 0.592 |
| BC | 0.654 | 0.654 | 0.644 | 0.644 | 0.630 | 0.653 | 0.649 | 0.649 |
| IBD | 0.662 | 0.654 | 0.658 | 0.662 | 0.636 | 0.665 | 0.654 | 0.651 |
| T2D | 0.624 | 0.619 | 0.626 | 0.624 | 0.592 | 0.625 | 0.612 | 0.620 |
| SCZ | 0.684 | 0.673 | 0.686 | 0.679 | 0.664 | 0.681 | 0.680 | 0.672 |
| BP | 0.612 | 0.607 | 0.613 | 0.612 | 0.604 | 0.612 | 0.609 | 0.608 |

Supplementary Table 8. The mean of AUC across 10 random splits for six diseases.

| Traits | SDPR | PRS-CS auto | SBayesR | LDpred2 auto |
|---|---|---|---|---|
| Height | 0.271 | 0.271 | 0.226 | 0.224 |
| BMI | 0.093 | 0.089 | 0.082 | 0.070 |
| HDL | 0.116 | 0.114 | 0.033 | 0.063 |
| LDL | 0.147 | 0.141 | 0.026 | 0.113 |
| Total cholesterol | 0.146 | 0.144 | 0.032 | 0.074 |
| Triglycerides | 0.075 | 0.082 | 0.021 | 0.055 |

Supplementary Table 9. The mean of variance of phenotypes explained by PRS across 10 random splits for six quantitative traits for methods without the need of parameter tuning.

| Traits | SDPR | PRS-CS auto | SBayesR | LDpred2 auto |
|---|---|---|---|---|
| CAD | 0.591 | 0.594 | 0.579 | 0.585 |
| BC | 0.654 | 0.652 | 0.644 | 0.617 |
| IBD | 0.662 | 0.652 | 0.658 | 0.616 |
| T2D | 0.624 | 0.619 | 0.626 | 0.602 |
| SCZ | 0.684 | 0.677 | 0.686 | 0.677 |
| BP | 0.612 | 0.610 | 0.613 | 0.610 |

Supplementary Table 10. The mean of AUC across 10 random splits for six diseases for methods without the need of parameter tuning.

### Supplementary Figure 2

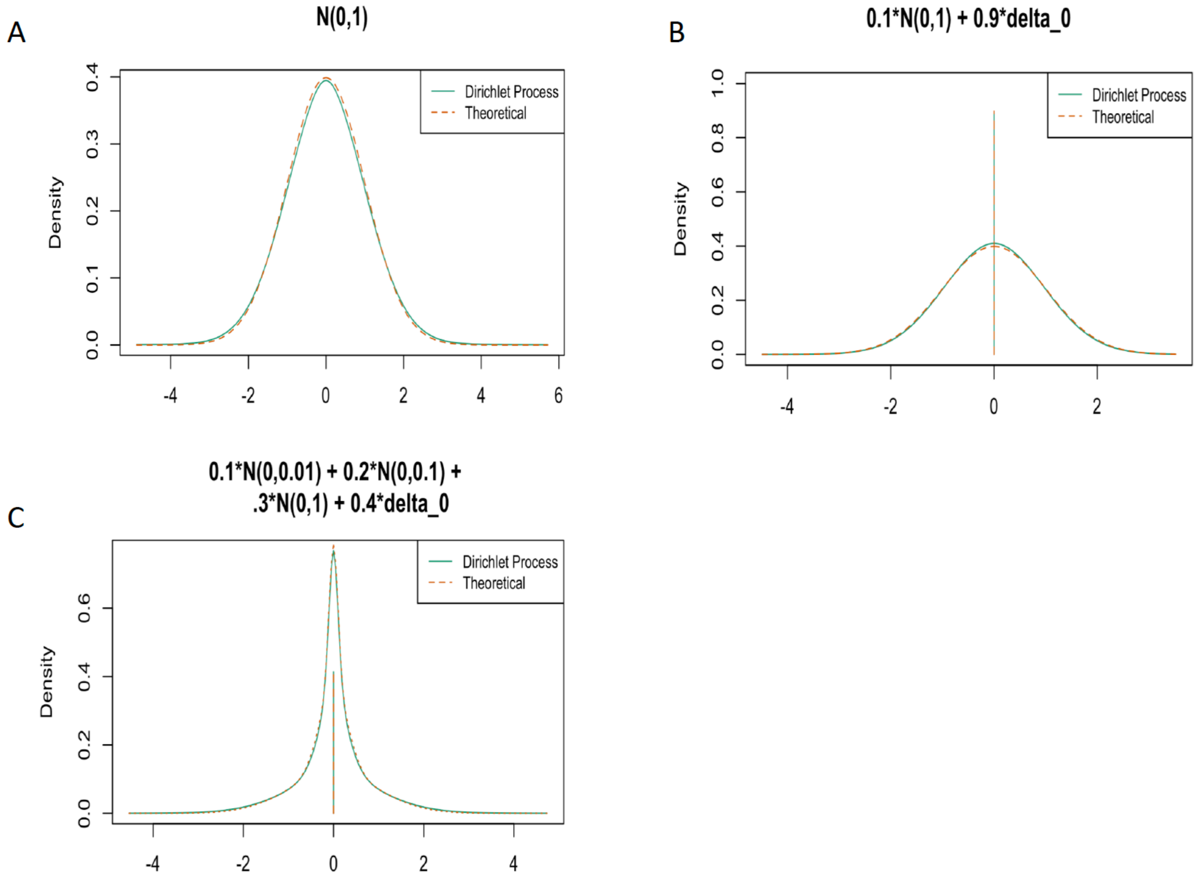


Supplementary Figure 2. Adaptiveness of Dirichlet process prior to different parametric assumptions. 1A: 2000 data were simulated from $N(0,1)$ satisfying the assumption of LDpred-inf. 1B:2000 data were simulated from $0.1N(0,1) + 0.9\delta_0$ satisfying the assumption of LDpred. 1C: 2000 data were simulated from $0.1N(0, 0.01) + 0.2N(0, 0.1) + 0.3N(0,1) + 0.4\delta_0$ satisfying the assumption of SBayesR. Theoretical and Dirichlet process fitted density was shown in the plot.

**Supplementary Figure 3**

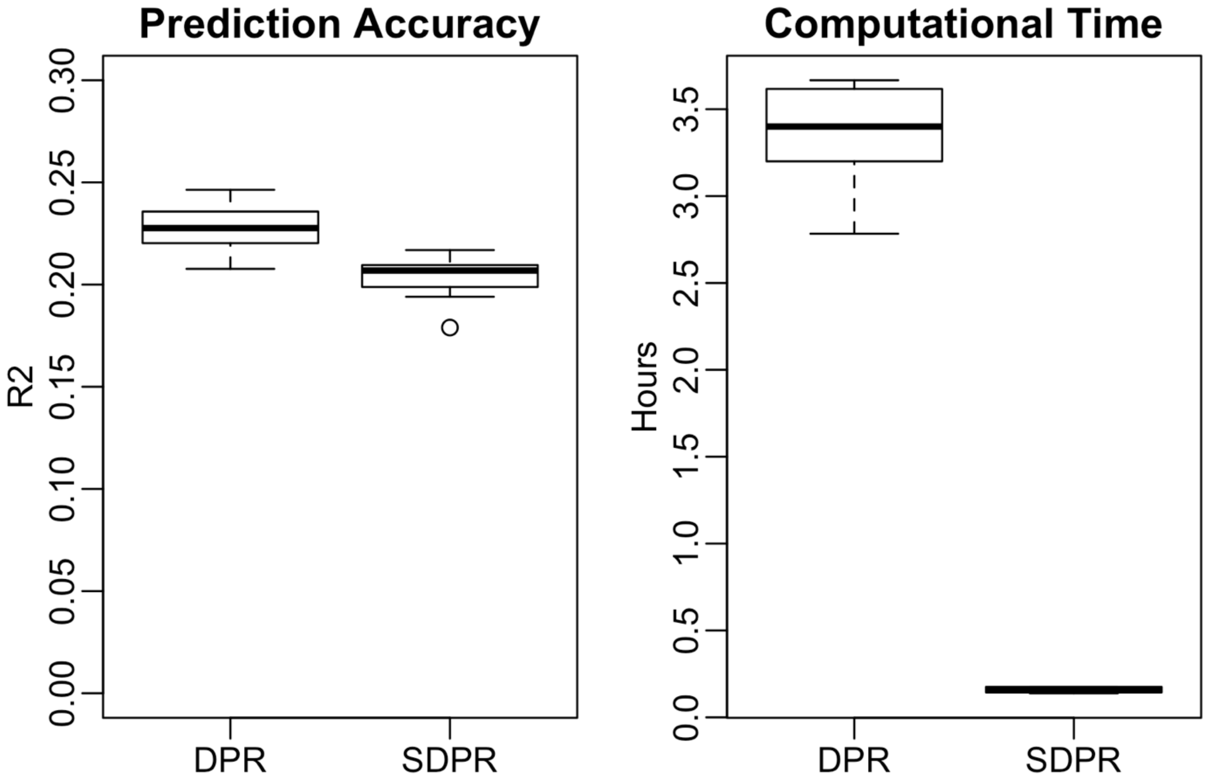


Supplementary Figure 3. Performance and Computational time of SDPR with DPR under a small-scale simulation using 10,000 individuals and 58,432 SNPs on chromosome 1. Effect sizes were generated as $\beta_j \sim \sum_{i=1}^{3} \pi_i N(0, c_i\sigma^2) + (1 - \sum_{i=1}^{3} \pi_i)\delta_0$ where $c = (1, 0.1, 0.01),\ \pi = (10^{-4}, 10^{-4}, 10^{-2})$ with $\sigma^2$ calculated so that the total heritability equaled 0.3. DPR was fit with 4 normal components, 2000 burnin and 4000 sampling iterations. SDPR was fit with 1000 maximum components and 1000 iterations. Simulation in each scenario was repeated for 10 times.

### References


1. Wood AR, Esko T, Yang J, Vedantam S, Pers TH, Gustafsson S, et al. Defining the role of common variation in the genomic and biological architecture of adult human height. Nat Genet. 2014;46(11):1173-86. Epub 2014/10/06. doi: 10.1038/ng.3097. PubMed PMID: 25282103; PubMed Central PMCID: PMCPmc4250049.
2. Willer CJ, Schmidt EM, Sengupta S, Peloso GM, Gustafsson S, Kanoni S, et al. Discovery and refinement of loci associated with lipid levels. Nat Genet. 2013;45(11):1274-83. Epub 2013/10/08. doi: 10.1038/ng.2797. PubMed PMID: 24097068; PubMed Central PMCID: PMCPMC3838666.
3. Gelman A. Prior distributions for variance parameters in hierarchical models (comment on article by Browne and Draper). Bayesian Anal. 2006;1(3):515-34. doi: 10.1214/06-BA117A.
4. Bulik-Sullivan BK, Loh PR, Finucane HK, Ripke S, Yang J, Patterson N, et al. LD Score regression distinguishes confounding from polygenicity in genome-wide association studies. Nat Genet. 2015;47(3):291-5. Epub 2015/02/03. doi: 10.1038/ng.3211. PubMed PMID: 25642630; PubMed Central PMCID: PMCPmc4495769.
5. Speed D, Balding DJ. SumHer better estimates the SNP heritability of complex traits from summary statistics. Nat Genet. 2019;51(2):277-84. Epub 2018/12/05. doi: 10.1038/s41588-018-0279-5. PubMed PMID: 30510236; PubMed Central PMCID: PMCPMC6485398.
6. Berisa T, Pickrell JK. Approximately independent linkage disequilibrium blocks in human populations. Bioinformatics. 2016;32(2):283-5. Epub 2015/09/24. doi: 10.1093/bioinformatics/btv546. PubMed PMID: 26395773; PubMed Central PMCID: PMCPmc4731402.
7. Khera AV, Chaffin M, Aragam KG, Haas ME, Roselli C, Choi SH, et al. Genome-wide polygenic scores for common diseases identify individuals with risk equivalent to monogenic mutations. Nat Genet. 2018;50(9):1219-24. Epub 2018/08/15. doi: 10.1038/s41588-018-0183-z. PubMed PMID: 30104762; PubMed Central PMCID: PMCPmc6128408.
